# KDRD: assessing discovery replicability based on summary association statistics from a single study

**DOI:** 10.64898/2026.11.03.763861

**Authors:** Zeyu Jiao, Yixin Zhang, Jujiao Kang, Liang Ma, Yawen Yang, Wei Zhao, Jianfeng Feng, Yinglei Lai

## Abstract

Replicability is essential for reliable discoveries in large-scale association studies. However, conventional replicability assessments typically require independent datasets or repeated data splitting, which may be constrained by data availability, computational burden and reduced sample sizes. Here, we developed Kernel Density-based Replicability of Discovery (KDRD), a statistical framework for estimating the expected replicability of statistically significant associations using summary statistics from a single study. Combining Gaussian kernel representations of observed association statistics with a theoretical null component, KDRD estimates replicability between independent studies under comparable conditions and enables sample size calculations beyond those currently available. In random subsampling simulations involving proteomics-based association analyses of 45 brain-related traits in the UK Biobank, KDRD showed a close agreement with the Dice Similarity Coefficients (DSCs). When KDRD was applied to the full available data,33 of the 45 traits showed at least moderate replicability, including 19 with high replicability. In matched-sample comparisons, proteomic associations showed higher KDRD estimates than pQTL associations across all 45 traits, whereas metabolomic associations showed higher estimates than proteomic associations for 33 traits. Higher protein missingness, proportions below the limit of detection, proportions of low-abundance proteins and assay variability were associated with lower KDRD estimates. KDRD-based sample size calculations suggested that approximately 200,000 and 500,000 participants would be needed for all 45 traits to achieve moderate and high replicability, respectively. Collectively, these findings demonstrate that KDRD provides a practical framework for quantitative replicability assessment and sample size calculations.

## Introduction

Replicability is a fundamental requirement for establishing robust scientific discoveries^1^. It generally refers to the ability to obtain consistent results across independent studies that address the same scientific question using newly collected data under sufficiently comparable experimental and analytical conditions^1,2^. The issue of replicability has received increasing attention with the widespread use of large-scale high-throughput data in biomedical research, including genomic, proteomic, metabolomic and neuroimaging association studies^3–6^. In many large-scale association studies, large numbers of associations are routinely evaluated and statistical summaries such as *p*-values or *z*-scores are obtained for each association^7,8^. Although false discoveries can be reduced by multiple-testing corrections in these analyses, it is still not directly assessed whether the association results can be consistently replicated across independent studies^9^. Therefore, replicability represents a distinct aspect of association reliability beyond that addressed by multiple-testing correction. Quantitative assessment of replicability is thus necessary in large-scale association studies^10,11^.

Replicability can be evaluated based on different aspects of consistency across independent experiments. Previous approaches have assessed the consistency of association rankings or effect directions^11–13^. For statistically significant findings, a particularly important criterion is whether a finding identified in one experiment can be identified again in another independent experiment. This can commonly be evaluated by examining the overlap of statistically significant findings, and quantified using similarity measures such as the Dice Similarity Coefficient (DSC) or Jaccard coefficient^14–16^. These two replicability measures, together with approaches based on other aspects of consistency^11–13^, generally require statistical summaries (i.e., *p* values or *z*-scores) from at least two independent datasets/experiments. However, in many studies, only a single dataset could be generally available.

When replicability assessment is limited to a single dataset, a common practical strategy is to divide the original dataset into two non-overlapping subsets and then to independently repeat the association analysis in each subset (i.e. random subsampling-based replicability assessment) ^6,17,18^. This provides an empirical assessment of internal replicability^6,17,18^. However, there is a clear limitation: each subset contains only a fraction of the original dataset. Because the statistical significance and the stability of association discoveries are generally influenced by the related sample size^6,19–21^, a replicability assessment based on reduced subsets may differ substantially from that based on the orignal/complete dataset. Therefore, this limitation can impact the interpretation of replicability.

In addition, the applicability of random subsampling-based replicability assessment may be limited by the availability of public data^22,23^. Individual-level data from large-scale biomedical studies may be subject to access restrictions^24,25^. This can make it difficult to perform random subsampling of the original dataset. Moreover, the applicability of random subsampling-based replicability assessment may also be limited by the burden of computing. As sample size and data dimensionality increase, repeatedly splitting individual-level data and rerunning association analyses impose greater computational and storage requirements^26,27^, which can be difficult to be achieved in practice. Therefore, these computational and practical limitations can restrict random subsampling-based replicability assessments in large-scale high-throughput studies.

When an independent external dataset is available, external validation provides an alternative to data splitting (as aboved described). However, heterogeneity in experiment conditions between the original and external datasets may introduce additional variability into replicability assessment^4,28,29^. In general, a replication study should address the same scientific question under sufficiently comparable study and analytical conditions, including population characteristics, phenotype definitions, molecular measurements, data processing and statistical procedures^4,28,29^. Importantly, it has been shown in a previous study that, even when the same neuroimaging dataset and hypotheses are used, differences in analytical workflows can lead to substantially different results^30^. This highlights the importance of comparability in study design as well as analytical procedures when evaluating replicability. Accordingly, heterogeneity in these conditions between the original and external datasets may contribute additional variability to replicability assessments. This issue is particularly relevant for large-scale biomedical resources, such as the UK Biobank, for which independent datasets with closely comparable study and analytical conditions may be difficult to obtain^4,31,32^. Together, these limitations motivate the development of a more practical approach to replicability assessment based solely on the original study, without requiring repeated subsampling or a closely matched external dataset.

A theoretical framework linking sample size to replicability could provide a basis for evaluating replicability across different sample sizes, and determining the sample size required to achieve a desired level of replicability. Larger sample sizes generally yield more stable association estimates and more replicable findings^6,20,21^. However, increasing sample size often requires longer periods of participant recruitment and data collection, as well as greater financial costs^33,34^. Therefore, it is important to perform sample size calculations beforehand. Determining the desirable sample size to achieve a required level of replicability is essential in the design of large-scale association studies^35,36^. Existing sample size calculations have typically examined the relationship between sample size and replicability through random subsampling of available datasets, with replicability assessed across a range of reduced sample sizes^6,35^. Although informative, these empirical evaluations are inherently bounded by the total sample size of the orignal data, and cannot directly assess replicability at larger target sample sizes. With the development of a theoretical framework, this limitation can be addressed.

Here, we developed the Kernel Density-based Replicability of Discovery (KDRD), a statistical framework for estimating the expected replicability of statistically significant discoveries using statistical summries based on one dataset. KDRD estimates the expected proportion of significant associations that would be replicated in a hypothetical independent study conducted under identical conditions, without splitting the original data or requiring a closely matched external dataset. Our framework further incorporates sample-size calculations to assess replicability across different target sample sizes. We first evaluated the performance of KDRD using random subsampling analyses in the UK Biobank. These analyses covered forty five brain-related traits, including physical measures, cognitive function, mental health and brain imaging metrics. The resulting KDRD estimates were compared with Dice Similarity Coefficients (DSCs). Then, we applied KDRD to all available samples for each trait to assess the replicability of discoveries from proteomics-based association studies. Using the same framework, we further compared replicability between proteomic and genomic associations as well as between proteomic and metabolomic associations. We also investigated the relationships between proteomic data-quality factors and KDRD-estimated replicability. Finally, the sample-size calculation component of KDRD was used to estimate the sample sizes required to achieve predefined levels of replicability.

## Methods

### Study participants and multi-omics data processing

The UK Biobank (UKB) is a large prospective cohort comprising more than 500,000 participants recruited across the United Kingdom. The resource provides extensive phenotypic information together with biological samples, genetic data, and imaging measurements^37^. In this study, we used proteomic, genomic, metabolomic, imaging, and phenotypic data from the UKB. This research was conducted using the UK Biobank resource under application number 19542, and data use complied with the terms and conditions of the UK Biobank. This study was conducted in accordance with the principles of the Declaration of Helsinki. All participants provided explicit, written informed consent to the UKB. The UKB cohort received approval from the NHS National Research Ethics Service North West (reference number: 16/NW/0274).

Plasma proteomic measurements were obtained using the Olink Explore platform, with protein levels reported as normalized protein expression (NPX) values^38^. Samples and proteins with more than 60% missing proteomic measurements were excluded. For the remaining data, missing NPX values were imputed using the median within groups defined by sex and age quartiles. The processed proteomic dataset comprised 52,632 participants and 2,920 proteins.

Genomic data were obtained from the UKB version 3 imputed dataset. Participants and genetic variants that passed the UKB quality-control procedures were retained for analysis^39^. We focused on 6,386 protein quantitative trait locus (pQTL)-related variants identified in previous studies^38,40^. To ensure that genomic and proteomic association analyses were conducted in the same participants, the two datasets were matched at the individual level, resulting in 43,685 participants with both genomic and proteomic data. Plasma metabolomic data were generated using the Nightingale Health nuclear magnetic resonance (NMR) platform, and 249 metabolic biomarkers were included^41^. The metabolomic and proteomic datasets were similarly matched at the individual level by retaining participants with available measurements in both datasets, resulting in 30,079 participants for the metabolomics–proteomics analyses.

Brain imaging measures were derived from T1-weighted structural MRI data processed by UKB. In the present study, we included total cortical surface area (CSA), total cortical volume (CV), and mean cortical thickness (CT) for the left and right cerebral hemispheres^42^.

For each phenotype, analyses were restricted to participants with available measurements for the corresponding phenotype and molecular data. Age and sex were additionally required for proteomic and metabolomic analyses, whereas genetic analyses further required information on the first 20 genetic principal components. Participants with missing values for any variable included in the corresponding association model were excluded. Consequently, the final sample size varied across phenotypes and molecular data types. Further details on data acquisition, quality control, and preprocessing procedures have been described in our previous study^40^.

### Association analysis and calculation of z-scores

Associations between molecular features and phenotypes were evaluated using general linear models. Age and sex were included as covariates in the proteomic and metabolomic analyses, while the genetic analyses additionally adjusted for the first 20 genetic principal components.

To place association statistics from proteomic, genomic, and metabolomic analyses on a common scale, the upper-tail *P* value for each molecular feature–phenotype association was converted to a z-score according to

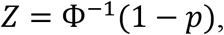

where *p* denotes the upper-tail *P* value and Φ^⁻1^ represents the inverse cumulative distribution function of the standard normal distribution. The resulting z-scores summarized the direction and statistical strength of the associations and were used in further analyses.

### Dice Similarity Coefficient (*DSC*)

A large number of z-scores (see above for its definition) *z* can be obtained from large-scale association analyses. With a threshold value *c*_+_ for the positive changes and a threshold value *c*_−_ for the negative changes, we can declare statistically significant results based on *z>c*_*+*_ or *z<c*_*-*_. In general, both threshold values are set the same as c.

Dice Similarity Coefficient (*DSC*) has been widely used to assess the replicability between two lists of statistically significant results from two association analyses.

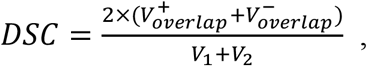

where *V*_1_ and *V*_2_ are the numbers of statistically significant results in the first and second analyses, respectively; *V*^**+**^ or *V*^**-**^ are the numbers of statistically significant results with *z>c*_*+*_ or *z<c*_*-*_ in both analyses.

According to this definition, *DSC* quantifies the proportion of statistically significant results that are replicated with the same direction across two experiments. Let *Z*^(1)^ represent the *z*-score from the original experiment and *Z*^(2)^ represent the corresponding *z*-score from an independent replication experiment for macthed study variable.Then, we can express the probabilistic definition for *DSC* as follows.

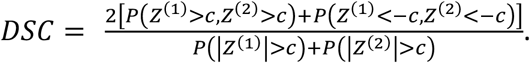

### Kernel Density-based Replicability of Discovery (*KDRD*)

Our objective was to assess the expected replicability of statistically significant results using only the association statistics obtained from one observed study. Let 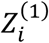 denote the z-score for the *i* th association unit obtained from a large-scale high-throughput association analysis in the observed study, and let 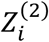 denote the corresponding z-score that would be obtained from an independent hypothetical replication study under comparable experimental conditions.

To distinguish the underlying association signal from study-specific sampling variation, we introduced a latent standardized effect *Θ*_*i*_. For study *r*, the association statistic was represented as

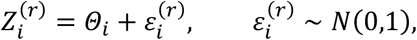

where *Θ*_*i*_ represents the underlying standardized association signal shared across comparable studies.

The sampling errors from the two independent studies were assumed to be conditionally independent given the latent effect, i.e.,

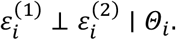

Thus, conditional on *Θ*_*i*_ = *θ*_*i*_,

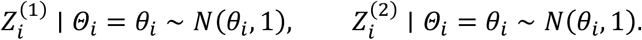

This formulation separates the latent association signal that is expected to persist across comparable studies from study-specific sampling variation. Across association units, the latent effects were represented using a two-group mixture distribution,

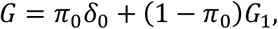

where *π*_*O*_ denotes the proportion of true null associations, *δ*_*O*_ denotes a point mass at zero, and *G*_1_ denotes the latent-effect distribution among non-null associations. Under the null hypothesis, *Θ* = 0, and therefore

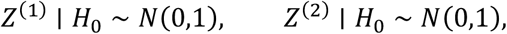

with conditional independence between the two experiments. The corresponding two-dimensional theoretical null density is

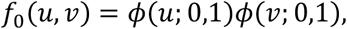

where *ϕ*(*x*; *μ, σ*^2^) denotes the probability density function of a normal distribution with mean *μ* and variance *σ*^2^, evaluated at *x*.

For a non-null association with latent effect *Θ* = *θ*, the joint density of the statistics from the original and hypothetical replication studies is

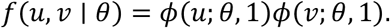

Integrating over the non-null latent-effect distribution gives

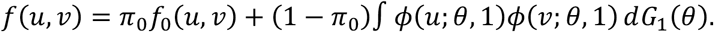

In practice, the latent non-null effect distribution *G*_1_ is unknown and is not directly observed from a single study. Let 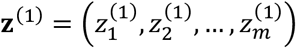 denote the observed z-scores from the reference study. Rather than attempting to directly estimate *G*_1_, we constructed a working empirical component based on the distribution of the observed z-scores as

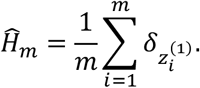

Here, the observed z-scores were used as empirical kernel locations to provide a data-driven representation of the association statistics in the reference study. This empirical component was used as a working plug-in representation for constructing the joint distribution, rather than as a direct estimate of the latent non-null effect distribution *G*_1_. Substituting this empirical approximation into the latent-effect model gives the empirical two-dimensional Gaussian-kernel component

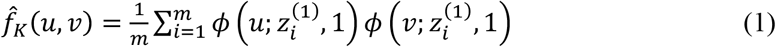

The proportion of true null hypotheses, *π*_*O*_, was estimated from the two-sided *P* values using the “convest” function implemented in the limma R package. The empirical Gaussian-kernel component in Eq. (1) and the theoretical null component were then combined using weights 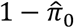 and 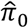, respectively,

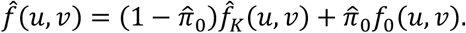

This formulation allows the probabilities required for the probabilistic DSC to be estimated without assigning a null or non-null status to each individual association.

Let 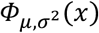 denote the cumulative distribution function of a normal distribution with mean *μ* and variance *σ*^2^, evaluated at *x*. Conditional on a latent effect *Θ* = *θ*, the probability of obtaining a statistically significant result in a single experiment at a two-sided z-score threshold *c* > 0 is

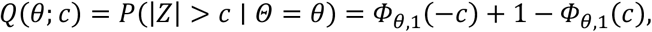

whereas the probability that the original and hypothetical replication experiments both yield statistically significant results in the same direction is

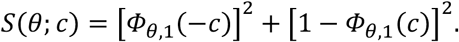

Accordingly, the population-level KDRD can be expressed as

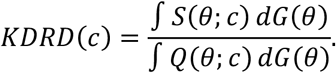

Under the empirical component described above, this population-level functional reduces to the computational form based on the observed z-scores. For the Gaussian component centered at 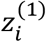, the probabilities of obtaining a negatively significant and positively significant result in a single experiment are, respectively,

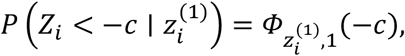

and

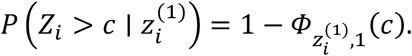

Because the original and hypothetical replication experiments are conditionally independent given the component location 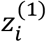 under the empirical component, the probability of obtaining a negatively significant result in both experiments is

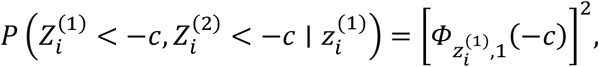

and the probability of obtaining a positively significant result in both experiments is

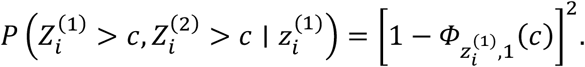

Therefore, the probability contributed by the *i*th empirical component to the numerator of the probabilistic *DSC* is

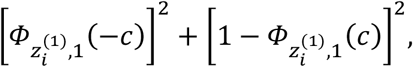

whereas its contribution to the denominator is the probability of obtaining a statistically significant result in a single experiment,

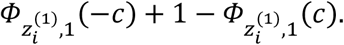

For the theoretical null component centered at zero, the corresponding numerator and denominator contributions are

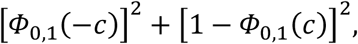

and

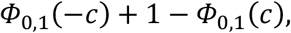

respectively. Combining the empirical Gaussian-kernel and theoretical null components using the estimated mixture weights, and summing over all observed z-scores, gives the Kernel Density-based Replicability of Discovery (*KDRD*):

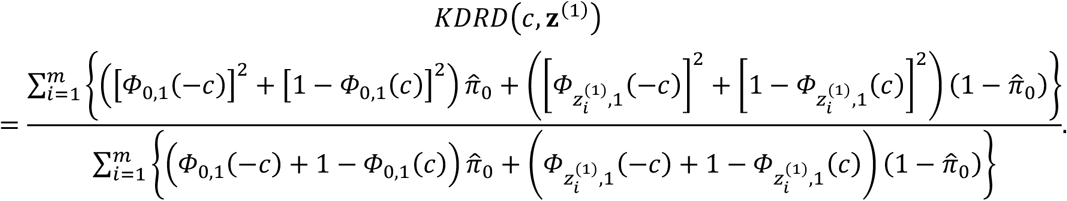

The numerator estimates the probability of obtaining statistically significant results with the same direction in both the original and hypothetical replication experiments, whereas the denominator estimates the probability of obtaining a statistically significant result in a single experiment. Thus, a larger *KDRD* value indicates a higher expected proportion of statistically significant discoveries that would remain statistically significant with the same direction in an independent hypothetical replication study conducted under comparable conditions. An overview of the analytical procedures for DSC and KDRD is provided in Extended Data Fig. 1.

### Sample size adjustment for *KDRD*

The latent-effect formulation allows KDRD to be evaluated under target sample sizes different from that of the observed reference study. Recall that Θ_*i*_ is defined as the latent standardized association effect, or equivalently the latent noncentrality parameter, at this reference information level. Under comparable study designs, let *β*_*i*_ denote the underlying association effect for the *i*th association unit, which is assumed to remain fixed across the reference and target studies. Suppose that the standard error of the corresponding effect estimator satisfies

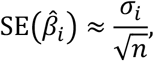

where *n* is the sample size of the observed reference study and *σ*_*i*_ denotes the asymptotic scale parameter determining the standard error of the effect estimator. The reference-study z-statistic can then be written approximately as

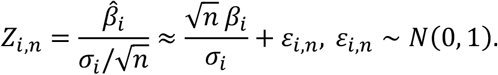

We therefore define the latent standardized association effect Θ_*i*_ at the reference information level as

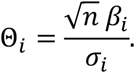

For a target study with sample size *m*_*r*_, the corresponding noncentrality parameter becomes

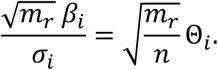

Accordingly, let 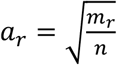, the statistic in target study *r* can be represented as

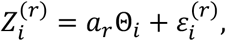

With 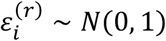.

The study-specific sampling errors are additionally assumed to be conditionally independent given the latent effect, i.e.,

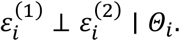

Therefore, conditional on Θ_*i*_ = *θ*_*i*_,

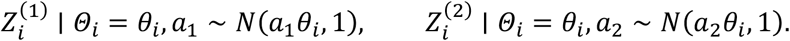

When the first target study is the observed reference study itself, *m*_1_ = *n* and hence *a*_1_ = 1, while *a*_2_ may be varied to evaluate the replicability expected for a replication study of a different size. Alternatively, both *m*_1_ and *m*_2_ may be varied when KDRD is used prospectively to compare designs for two future studies.

Under the empirical approximation 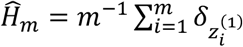, the latent component locations are represented computationally by the observed 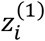. Consequently, the component means in the two target studies become 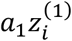 and 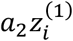, respectively, while the unit sampling variance is retained. Thus,

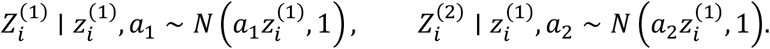

For the theoretical null component, the component mean remained zero regardless of sample size. When *a*_1_ ≠ *a*_2_, the two target studies no longer necessarily have the same marginal z-score distribution. The sample-size-adjusted *KDRD* was defined as

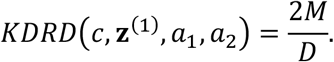

Further details on the definitions of *M* and *D* are provided below. Using the same *S* notation as above, we defined the same-direction significance probability for the theoretical null component as

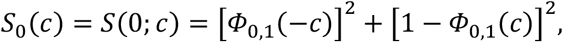

and, for the empirical component centered at 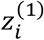, the corresponding probability after sample-size adjustment as

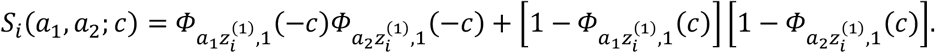

The numerator was then calculated as

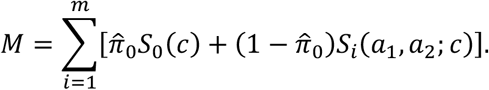

Using the same *Q* notation as above, we defined the probability of obtaining a statistically significant result from the theoretical null component as

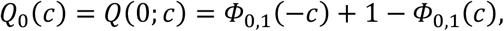

and the corresponding probability for the empirical component in target study *j* as

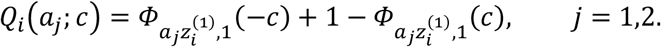

Because the denominator of the general probabilistic DSC is the sum of the significant-result probabilities in the two studies, it was calculated as

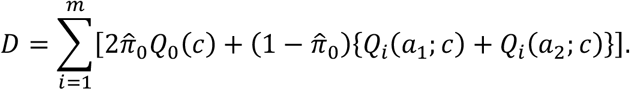

Here, 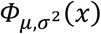 denotes the cumulative distribution function of a normal distribution with mean *μ* and variance *σ*^2^, evaluated at *x*, as defined above. The terms involving 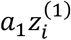 and 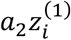 represent the sample-size-adjusted empirical Gaussian components for the two target studies, whereas the theoretical null component remains centered at zero. When *a*_1_ = *a*_2_ = 1, corresponding to target sample sizes equal to that of the reference dataset, this formulation reduces to the original KDRD calculation. This sample-size adjustment enables the expected replicability of statistically significant discoveries to be evaluated under different study sample sizes and can therefore be used to investigate the sample size required to achieve a desired level of discovery replicability.

In the practical application presented here, *N* denotes the nominal overall cohort size rather than the number of complete observations for each trait. The *N* = 50,000 setting represents the current dataset, for which KDRD was calculated using the actual available sample size for each trait. Projections for larger cohorts accounted for anticipated missingness when determining the trait-specific effective sample sizes used in the sample-size adjustment.

### Simulation analysis for KDRD

To evaluate the performance of KDRD, we conducted random subsampling simulations using 45 traits from four categories, including physical measures, cognitive function, mental health, and brain imaging metrics. For each trait, participants in the original dataset were randomly divided into two non-overlapping subsets of equal sample size, thereby generating two experiments with comparable study conditions. Association analyses were independently performed in the two subsets using the same analytical procedures, and the Dice Similarity Coefficient (DSC) was calculated between the two experiments to quantify the observed replicability of statistically significant findings. This random subsampling procedure was repeated 1,000 times for each trait.

In each random subsampling experiment, KDRD was calculated using only the association statistics obtained from the first subset, whereas DSC was calculated using the association results from both subsets. Thus, DSC served as an empirical reference for replicability based on two independently analyzed datasets, while KDRD estimated replicability using information from only one of the two datasets. For each trait, the 1,000 random subsampling experiments generated 1,000 DSC values and 1,000 corresponding KDRD estimates. The distributions of the two replicability measures were summarized by their median and interquartile range (Q1–Q3). The agreement between KDRD and DSC across traits was further evaluated using Spearman correlation analyses of their median, Q1, and Q3 values, respectively.

We further calculated the hitting rate (HR) to quantify whether the KDRD estimates were consistent with the empirical variation in DSC. For each trait, the 95% empirical interval of DSC was defined from the 1,000 DSC values, and HR was calculated as the proportion of the 1,000 KDRD estimates that fell within this interval. Specifically, if *DSC*_2.5%_ and *DSC*_97.5%_ denote the 2.5th and 97.5th percentiles of the 1,000 DSC values, respectively, HR was defined as

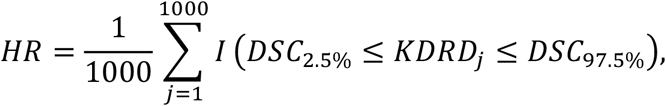

where *I*(⋅) is the indicator function and *KDRD*_*j*_ denotes the KDRD estimate from the *j*th random subsampling experiment. A higher HR therefore indicates that a larger proportion of KDRD estimates obtained from a single subset were within the empirical 95% variation range of DSC derived from two comparable experiments.

### Relationships between proteomic data-quality factors and KDRD

To quantitatively assess the relationships between proteomic data-quality factors and KDRD, we performed a resampling-based analysis that generated protein sets with systematically varying data-quality profiles. Specifically, 300 proteins, corresponding to approximately 10% of the available proteins, were randomly sampled for each analysis. Proteins were drawn from the UK Biobank “Explore 1,536 assay panels” and “Expansion 1460 assay panels”, hereafter referred to as Panel 1 and Panel 2, respectively. To generate protein sets with different panel compositions, the proportion of proteins sampled from Panel 1 was varied from 0% to 99% in 1% increments, with the remaining proteins sampled from Panel 2. For each proportion, the sampling procedure was independently repeated 10 times, resulting in a total of 1,000 sampled protein sets.

For each sampled protein set, we calculated the mean protein-level missing rate, mean proportion of samples below the limit of detection (LOD), mean protein-level coefficient of variation (CoV), and the proportion of low-abundance proteins across the 300 proteins. Low-abundance proteins were defined as proteins measured at a dilution level of 1:1, and their proportion was calculated as the number of such proteins divided by the total number of proteins in each sampled set. Association analyses were then performed for each sampled protein set, and the corresponding KDRD was calculated using a Bonferroni-corrected discovery threshold of *P* < 0.05/300 (approximately 1.67 × 10^−4^). Spearman correlation analysis was used to evaluate the relationships between each proteomic data-quality factor and KDRD. This resampling framework generated a broad range of proteomic data-quality profiles while maintaining a fixed number of proteins across analyses, allowing the relationships between these factors and KDRD to be quantitatively evaluated.

## Results

### Motivation

In many studies, the original individual-level data are not publicly available and may be subject to access restrictions, posing a practical barrier to replicability assessment based on random subsampling of the original dataset. The computational burden of random subsampling-based replicability assessment is also an important practical consideration in large-scale omics studies. With increasing sample sizes and numbers of measured features, repeated subsampling experiments and corresponding association analyses impose considerable computational and storage requirements. These practical constraints can therefore hinder the application of random subsampling-based replicability assessment in large-scale omics association studies.

Even when the original data are accessible and random subsampling is computationally feasible, internal replicability assessment is performed based on reduced sample size. To investigate how this reduction affects the replicability assessment, we conducted random subsampling experiments across 45 brain-related traits. For each trait, participants were randomly divided into two non-overlapping subsets, with each subset containing 1/8, 1/4, or 1/2 of the total available sample for that trait. Association analyses were performed independently in the two subsets using the same analytical procedure, and the overlap of statistically significant associations with consistent directions was quantified using the Dice Similarity Coefficient (*DSC*). Statistical significance was defined using a Bonferroni-corrected threshold of *P* < 1.7 × 10^−5^, corresponding to 0.05/2,920 tested proteins. For each sample-size setting, the random subsampling procedure was repeated 1,000 times, and the median *DSC* with interquartile range (Q1–Q3) across the 1,000 paired experiments was used to summarize replicability.

A clear decrease in *DSC* was observed as the sample size decreased (Fig. 1). At the 1/8 sample size, only 2 of the 45 phenotypes showed moderate replicability (0.5 ≤ *DSC* < 0.75), and 2 showed high replicability (*DSC* ≥ 0.75). At the 1/4 sample size, 8 traits showed moderate replicability and 4 showed high replicability. When the sample size increased to 1/2, 9 traits showed moderate replicability and 12 showed high replicability. Accordingly, the number of traits with moderate-to-high replicability increased from 4 at the 1/8 sample size to 12 at the 1/4 sample size and 21 at the 1/2 sample size. Detailed DSC results for each trait and sample-size setting are provided in Supplementary Table 1. These results confirm that *DSC*-based internal replicability assessment can be markedly affected by reductions in the sample size of the paired experiments.

**Fig. 1.**
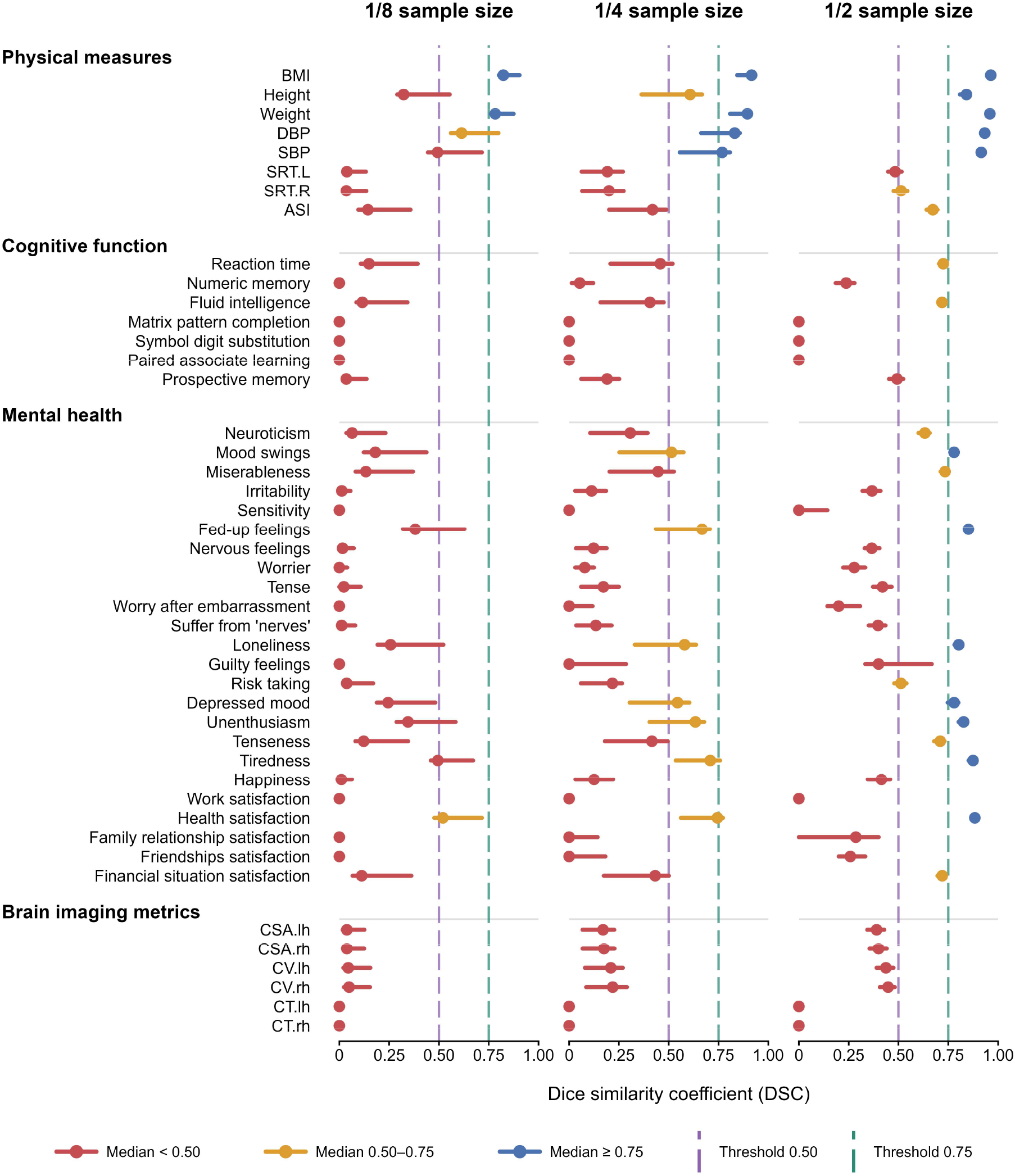
Effect of sample size on DSC-based replicability. For each of the 45 phenotypes, participants were randomly divided into two non-overlapping subsets, each containing 1/8, 1/4, or 1/2 of the phenotype-specific total sample size. The random subsampling procedure was repeated 1,000 times. Points indicate the median *DSC* and horizontal lines indicate the interquartile range (Q1–Q3). Dashed vertical lines indicate DSC thresholds of 0.50 and 0.75.

External validation provides an alternative strategy for replicability assessment without splitting the original dataset. However, differences in study and analytical conditions between the original and external datasets may introduce additional variability into the assessment. A suitable replication dataset should address the same scientific question under sufficiently comparable conditions, including population characteristics, phenotype definitions, molecular measurements, data processing and statistical procedures. For large-scale biomedical resources such as the UK Biobank, independent datasets that are closely comparable across these dimensions may be difficult to obtain. This limits the practical applicability of external validation for replicability assessment.

Together, these limitations motivated the development of a more practical approach to replicability assessment based solely on the original data, without the need for repeated subsampling or a closely matched external dataset.

### Simulation results

We next evaluated the performance of KDRD using random subsampling simulations across 45 brain-related traits, including physical measures, cognitive function, mental health, and brain imaging metrics (see the “Simulation analysis for KDRD” section in Methods). Both DSC and KDRD were calculated using the same Bonferroni-corrected significance threshold of *P* < 1.7 × 10^−5^ (0.05/2920). Overall, KDRD showed a high degree of agreement with the empirically derived DSC across the 45 traits (Fig. 2a). The median absolute difference between the median DSC and KDRD was 0.018, and 33 of the 45 traits showed an absolute difference of less than 0.05.

**Fig. 2.**
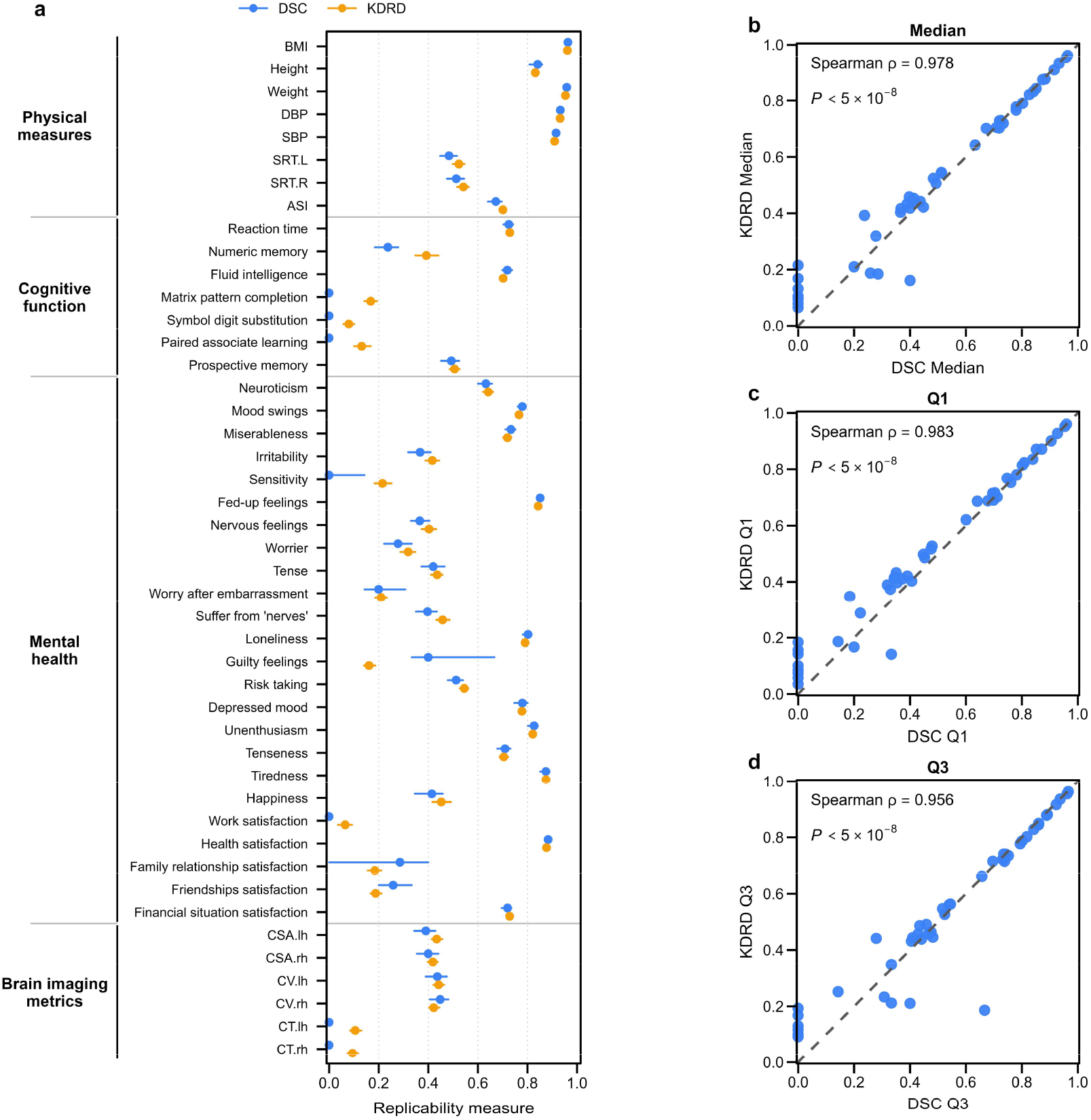
Comparison of DSC and KDRD across random subsampling simulations. **a**, Median DSC and KDRD across 1,000 simulations for each of the 45 traits; horizontal lines indicate Q1–Q3. **b–d**, Correlations between DSC and KDRD for the median (**b**), Q1 (**c**), and Q3 (**d**) across traits. Both measures were calculated using a Bonferroni-corrected significance threshold of *P* < 1.7 × 10^−5^. Spearman correlation coefficients and corresponding *P* values are shown.

This agreement was further supported by the distributions obtained from the 1,000 random subsampling simulations. Across traits, the median KDRD was strongly correlated with the median DSC (Spearman *ρ* = 0.978, *P* < 5 × 10^−8^; Fig. 2b). Similarly strong correlations were observed for Q1 (Spearman *ρ* = 0.983, *P* < 5 × 10^−8^; Fig. 2c) and Q3 (Spearman *ρ* = 0.956, *P* < 5 × 10^−8^; Fig. 2d). These results indicate that KDRD captured not only the overall level of DSC-based replicability across traits, but also the variation in replicability observed across random subsampling simulations.

We further assessed the agreement between KDRD estimates and the empirical variation in DSC using the hitting rate (HR; see the “Simulation analysis for KDRD” section in Methods). Overall, HR was high across the 45 traits, with a median of 0.998 (Q1–Q3, 0.960–1.000). A total of 42 of the 45 traits had an HR of at least 0.90, 36 had an HR of at least 0.95, and 19 had an HR of 1.00. Trait-specific DSC, KDRD and HR results are provided in Supplementary Table 2. Together, these results support the ability of KDRD, calculated using association statistics from only one of the two subsampled datasets, to approximate the replicability empirically observed between two studies conducted under comparable conditions.

### Replicability of PBAS discoveries across brain-related traits

Using all available samples for each trait, we applied KDRD to characterize the replicability of discoveries from proteomics-based association studies (PBAS) across 45 brain-related traits, including physical measures, cognitive function, mental health, and brain imaging metrics. The available sample sizes ranged from 4,631 to 52,444 participants. KDRD was estimated using a Bonferroni-corrected significance threshold of *P* < 1.7 × 10^−5^ (0.05/2,920). Overall, KDRD estimates ranged from 0.090 to 0.970. Among the 45 traits, 19 (42.2%) showed high replicability (KDRD > 0.75), 14 (31.1%) showed moderate replicability (0.5 < KDRD ≤ 0.75), and 12 (26.7%) showed low replicability (KDRD ≤ 0.5). Thus, 33 of the 45 traits (73.3%) showed at least moderate replicability under their currently available sample sizes (Table 1).

**Table 1.** KDRD estimates for PBAS discoveries across brain-related traits.

| UK Biobank category | Trait | Sample size | KDRD |
|---|---|---|---|
| **Physical measures** | BMI | 52,374 | 0.970 |
|  | Height | 52,444 | 0.907 |
|  | Weight | 52,414 | 0.965 |
|  | DBP | 49,335 | 0.959 |
|  | SBP | 49,335 | 0.948 |
|  | SRT.L | 16,664 | 0.670 |
|  | SRT.R | 16,645 | 0.686 |
|  | ASI | 17,465 | 0.794 |
| **Cognitive function** | Reaction time | 51,987 | 0.806 |
|  | Numeric memory | 5,517 | 0.619 |
|  | Fluid intelligence | 16,886 | 0.797 |
|  | Matrix pattern completion | 4,631 | 0.312 |
|  | Symbol digit substitution | 4,637 | 0.090 |
|  | Paired associate learning | 4,679 | 0.255 |
|  | Prospective memory | 16,624 | 0.654 |
| **Mental health** | Neuroticism | 41,820 | 0.768 |
|  | Mood swings | 51,066 | 0.836 |
|  | Miserableness | 51,539 | 0.794 |
|  | Irritability | 50,046 | 0.598 |
|  | Sensitivity | 50,913 | 0.390 |
|  | Fed-up feelings | 51,371 | 0.885 |
|  | Nervous feelings | 51,132 | 0.546 |
|  | Worrier | 51,100 | 0.465 |
|  | Tense | 50,494 | 0.578 |
|  | Worry after embarrassment | 50,279 | 0.338 |
|  | Suffer from 'nerves' | 50,503 | 0.630 |
|  | Loneliness | 51,539 | 0.841 |
|  | Guilty feelings | 51,044 | 0.262 |
|  | Risk taking | 50,509 | 0.692 |
|  | Depressed mood | 50,131 | 0.839 |
|  | Unenthusiasm | 50,582 | 0.869 |
|  | Tenseness | 50,280 | 0.786 |
|  | Tiredness | 50,810 | 0.925 |
|  | Happiness | 17,576 | 0.632 |
|  | Work satisfaction | 12,153 | 0.136 |
|  | Health satisfaction | 17,594 | 0.905 |
|  | Family relationship satisfaction | 17,518 | 0.310 |
|  | Friendships satisfaction | 17,430 | 0.300 |
|  | Financial situation satisfaction | 17,513 | 0.824 |
| **Brain imaging metrics** | CSA.lh | 5,623 | 0.602 |
|  | CSA.rh | 5,623 | 0.578 |
|  | CV.lh | 5,623 | 0.577 |
|  | CV.rh | 5,623 | 0.546 |
|  | CT.lh | 5,623 | 0.199 |
|  | CT.rh | 5,623 | 0.163 |

PBAS, proteomics-based association studies. KDRD was estimated using all available samples for each trait. The discovery threshold was *P* < 1.7 × 10<sup>-5</sup> (Bonferroni correction, 0.05/2,920).

Replicability varied across trait categories. Physical measures generally showed high replicability: all eight traits showed at least moderate replicability, and six showed high replicability. BMI, weight, diastolic blood pressure, systolic blood pressure, and height had KDRD values of 0.970, 0.965, 0.959, 0.948, and 0.907, respectively. Cognitive traits showed greater variation, with KDRD ranging from 0.090 for symbol digit substitution to 0.806 for reaction time. Reaction time and fluid intelligence showed high replicability, whereas numeric memory and prospective memory showed moderate replicability.

Among the 24 mental health traits, 17 showed at least moderate replicability, including 11 with high replicability. Relatively high KDRD estimates were observed for tiredness (0.925), health satisfaction (0.905), fed-up feelings (0.885), unenthusiasm (0.869), loneliness (0.841), and depressed mood (0.839). In contrast, several traits showed relatively low replicability, including work satisfaction (0.136), guilty feelings (0.262), friendships satisfaction (0.300), family relationship satisfaction (0.310), and worry after embarrassment (0.338).

For brain imaging metrics, cortical surface area and cortical volume showed moderate replicability, with KDRD values ranging from 0.546 to 0.602 across the two hemispheres. In contrast, cortical thickness showed lower KDRD values in both the left (0.199) and right (0.163) hemispheres. Together, these results demonstrate considerable variation in the estimated replicability of PBAS discoveries across brain-related traits and indicate that, under the currently available sample sizes, most of the evaluated traits showed moderate-to-high replicability.

### Cross-omics comparison of association replicability based on KDRD

We next used KDRD to compare association replicability across different molecular layers. To ensure comparability within each analysis, comparisons were performed using matched samples between pQTLs and proteomics (*N* = 43,685) and between metabolomics and proteomics (*N* = 30,079) across the same 45 brain-related traits, including physical measures, cognitive function, mental health, and brain imaging metrics (details see “Study participants and multi-omics data processing” section in Methods). KDRD was estimated using omics-specific Bonferroni-corrected significance thresholds of *P* < 1.7 × 10^−5^ for proteomics (0.05/2,920), *P* < 2.0 × 10^−4^ for metabolomics (0.05/249), and *P* < 7.8 × 10^−6^ for pQTLs (0.05/6,386).

In the matched comparison between pQTLs and proteomics, proteomics showed higher KDRD estimates for all 45 traits (Fig. 3). The median KDRD across traits was 0.641 for proteomics, compared with 0.055 for pQTLs. This pattern was consistently observed across all four trait categories, with proteomics showing higher KDRD estimates for all 8 physical measures, 7 cognitive function traits, 24 mental health traits, and 6 brain imaging metrics. The differences were pronounced for a number of traits. For example, KDRD estimates for health satisfaction were 0.895 for proteomics and 0.035 for pQTLs, while those for BMI were 0.968 and 0.118, respectively.

**Fig. 3.**
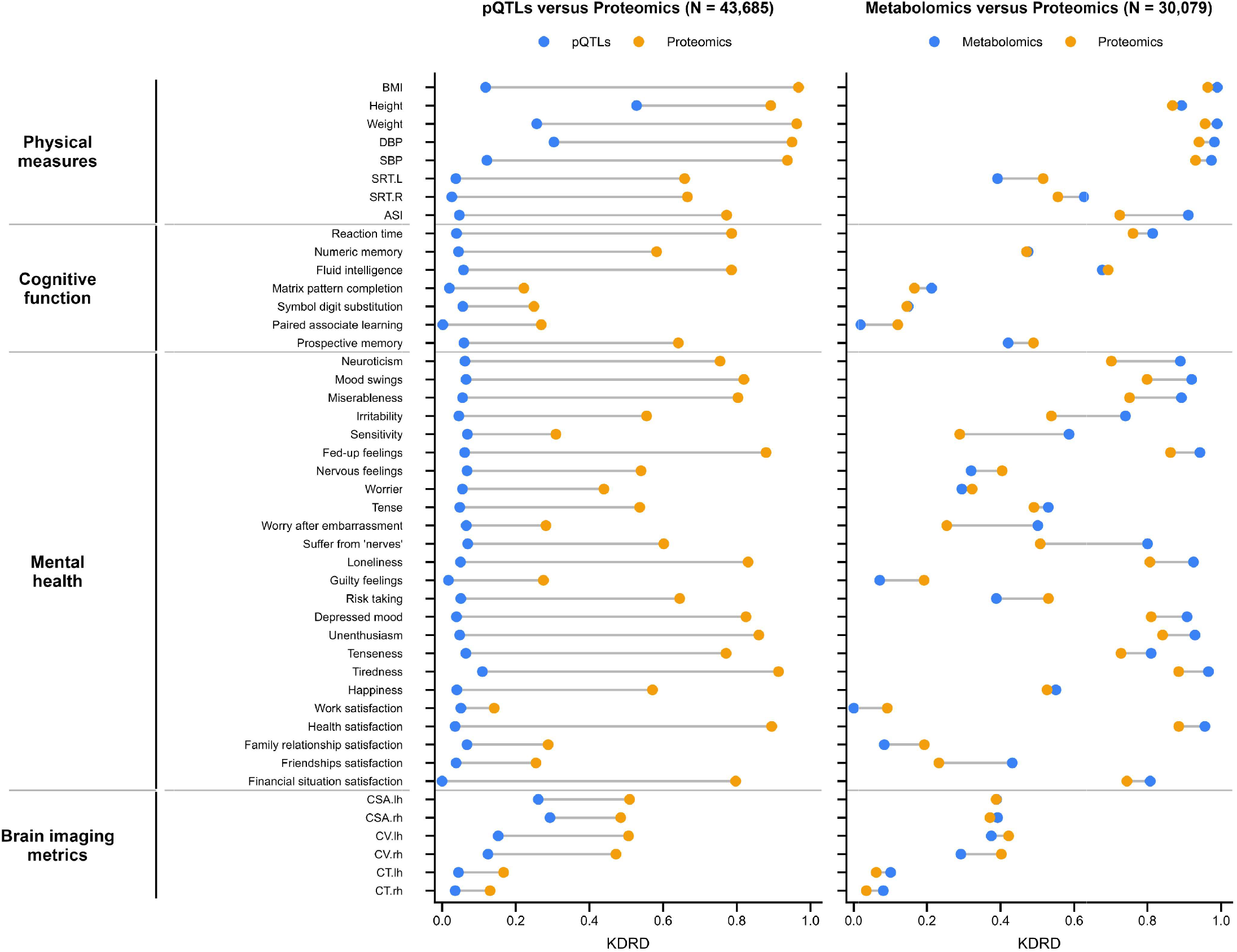
KDRD assessments across different molecular layers. KDRD assessments were compared between pQTLs vs. proteomics (*N* = 43,685; left) and between metabolomics vs. proteomics (*N* = 30,079; right) across 45 brain-related traits. Paired points represent the same trait across different molecular layers.

The comparison between metabolomics and proteomics showed a less uniform pattern. The median KDRD across traits was 0.586 for metabolomics and 0.526 for proteomics. Metabolomics showed higher KDRD estimates for 33 of the 45 traits (73.3%), whereas proteomics showed higher estimates for the remaining 12 traits. This pattern differed across trait categories: metabolomics showed higher KDRD estimates for 7 of 8 physical measures, 4 of 7 cognitive function traits, 18 of 24 mental health traits, and 4 of 6 brain imaging metrics. Several relatively large differences in favor of metabolomics were observed for sensitivity (0.586 versus 0.289), suffer from ‘nerves’ (0.801 versus 0.508), worry after embarrassment (0.501 versus 0.254), and irritability (0.741 versus 0.538). In contrast, proteomics showed higher KDRD estimates for several traits, including risk taking (0.530 versus 0.388), SRT.L (0.516 versus 0.392), guilty feelings (0.192 versus 0.072), and right-hemisphere cortical volume (0.402 versus 0.292). Trait-specific KDRD estimates for both matched-omics comparisons are provided in Supplementary Table 3. Overall, these matched-sample comparisons revealed marked differences in association replicability across molecular layers, with proteomics showing consistently higher KDRD than pQTLs, whereas the relative replicability of metabolomics and proteomics varied across traits.

### Proteomic data-quality factors were associated with KDRD

To illustrate the relationships between proteomic data-quality factors and KDRD, we present resampling results for two representative brain-related traits, fluid intelligence and neuroticism (see “Relationships between proteomic data-quality factors and KDRD” in Methods for details). For each sampled 300 protein set, we calculated *KDRD* together with the mean protein missing rate, mean proportion of samples below the LOD, proportion of low-abundance proteins and mean protein CoV (Fig. 4). For fluid intelligence, *KDRD* was negatively correlated with the mean protein missing rate (Spearman’s *ρ* = −0.73), mean proportion of samples below the LOD (*ρ* = −0.73), proportion of low-abundance proteins (*ρ* = −0.71) and mean protein CoV (*ρ* = −0.53; all *P* < 5 × 10^−8^). Similar patterns were observed for neuroticism, with corresponding Spearman correlations of −0.80, −0.79, −0.76 and −0.60, respectively (all *P* < 5 × 10^−8^). Together, these findings suggest that increases in protein missingness, the proportion of samples below the LOD, the proportion of low-abundance proteins and protein CoV may contribute to reductions in *KDRD*.

**Fig. 4.**
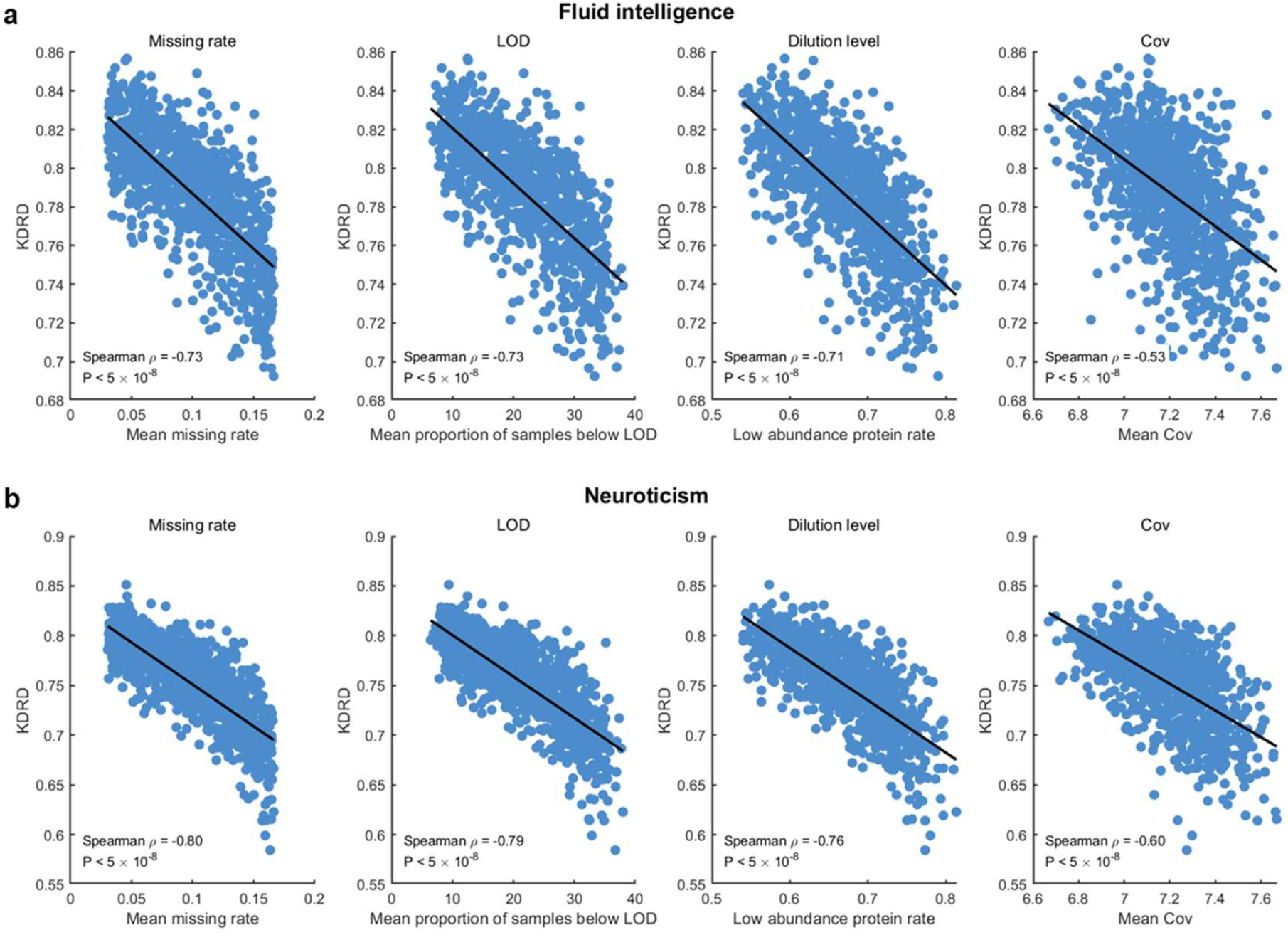
Relationships between proteomic data-quality factors and KDRD. Relationships of the mean protein missing rate, mean proportion of samples below the limit of detection (LOD), proportion of low-abundance proteins and mean protein coefficient of variation (CoV) with *KDRD* for (**a**) fluid intelligence and (**b**) neuroticism. Each point represents one sampled protein set. Black lines indicate linear fits for visualization. Spearman’s correlation coefficients (*ρ*) and corresponding *P* values are shown in each subfigure.

### KDRD-based sample size evaluation

To estimate the sample sizes required to achieve reliable replicability, we calculated *KDRD* for 45 brain-related traits at target sample sizes of 50,000, 100,000, 200,000 and 500,000 using the full dataset (Fig. 5). Details of the sample size evaluation are provided in the “Sample size adjustment for *KDRD*” section of the Methods. The number of traits reaching moderate or high replicability increased progressively with sample size. At a sample size of 50,000, 14 traits showed moderate replicability (0.5 < KDRD ≤ 0.75) and 19 showed high replicability (KDRD > 0.75), whereas 12 remained below the moderate-replicability threshold (KDRD < 0.5). At 100,000, 13 traits showed moderate replicability and 26 showed high replicability, leaving only six traits below the moderate threshold. When the sample size increased to 200,000, all 45 traits reached at least moderate replicability, including 33 traits with high replicability. At 500,000, all 45 traits reached high replicability.

**Fig. 5.**
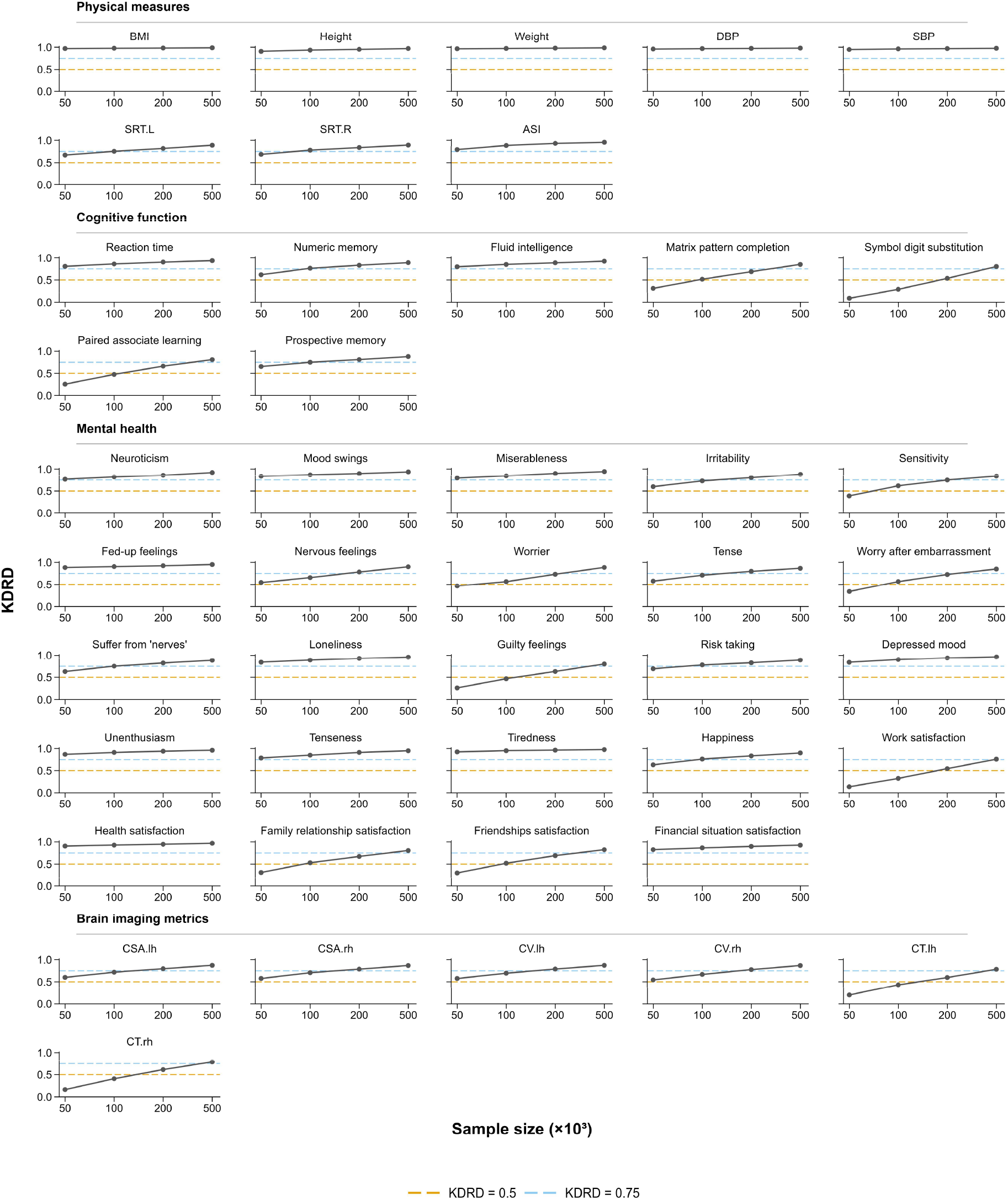
KDRD-based sample size evaluation across brain-related traits. *KDRD* values were estimated for 45 brain-related traits at target sample sizes of 50,000, 100,000, 200,000 and 500,000. Traits were grouped into physical measures, cognitive function, mental health and brain imaging metrics. Horizontal dashed lines indicate the thresholds for moderate (*KDRD* = 0.5) and high (*KDRD* = 0.75) replicability. CSA, cortical surface area; CV, cortical volume; CT, cortical thickness; lh, left hemisphere; rh, right hemisphere. Sample sizes indicate nominal overall cohort sizes; trait-specific effective sample sizes account for current or anticipated data missingness.

The required sample size also varied across trait domains. Physical measures generally achieved high replicability at relatively smaller sample sizes, whereas cognitive, mental health and brain imaging traits required larger sample sizes to reach comparable levels of replicability. In particular, several cognitive and brain imaging traits remained below moderate or high replicability at 50,000 - 100,000 samples but showed clear improvements at 200,000 and 500,000 samples. Projected KDRD values for each trait at the four target sample sizes are provided in Supplementary Table 4. Overall, these results suggest that a sample size of approximately 200,000 may be sufficient to achieve at least moderate replicability across the brain-related traits evaluated here. A sample size of 500,000 was sufficient to achieve consistently high replicability across all evaluated brain-related traits.

## Discussion

In this study, we developed KDRD, a statistical framework for estimating the expected replicability of statistically significant discoveries using association statistics from a single dataset. Unlike conventional overlap-based assessments that require two datasets or experiments, KDRD estimates the proportion of significant associations expected to be replicated in a hypothetical independent study conducted under comparable conditions. The framework therefore avoids the reduction in sample size introduced by data splitting and does not require a closely matched external dataset. Random subsampling analyses across 45 brain-related traits showed close agreement between KDRD and observed corresponding DSC, supporting the ability of KDRD to approximate the replicability observed between two independently analyzed samples. Using KDRD, we further characterized the replicability of proteomic association discoveries, compared replicability across molecular layers, examined its relationships with proteomic data-quality factors, and evaluated replicability across different target sample sizes.

An important motivation for KDRD is the dependence of conventional internal replicability assessment on data splitting. When a dataset is divided into independent subsets, the association analyses are necessarily performed with fewer samples than are available in the original study. Our results showed that DSC decreased markedly as the sample size of the paired experiments was reduced, indicating that replicability estimated from split samples can be strongly affected by the sample size used for assessment. KDRD addresses this limitation by estimating discovery replicability directly from the association statistics of a single dataset, without requiring data splitting. In the random subsampling simulations, KDRD assessments closely approximated the DSC obtained from paired subsets across diverse brain-related traits. For each trait, the median and interquartile range of KDRD and DSC were calculated across 1,000 simulations. Across the 45 traits, strong correlations were observed between KDRD and DSC for their median, first-quartile and third-quartile values. These findings suggest that the distribution of association statistics from a single study can provide an informative estimate of the expected replicability of association discoveries across comparable studies.

Applications of KDRD to proteomics-based association studies revealed substantial variations in discovery replicability across brain-related traits. Using all available samples for each trait, most traits showed at least moderate replicability, although clear differences were observed across trait categories. Physical measures generally showed high replicability, whereas cognitive function and mental health traits were more heterogeneous. Among brain imaging metrics, cortical surface area and cortical volume showed moderate replicability, whereas cortical thickness showed considerably lower values. Physical measures are generally assessed using relatively standardized and objective procedures, which may contribute to more stable association signals^37^. In contrast, cognitive function and mental health traits often involve greater heterogeneity in measurement and underlying biological architecture^29,43^, which may contribute to greater variations in association replicability. For brain imaging traits, previous studies have shown that cortical thickness generally has lower heritability, greater sensitivity to environmental influences and lower test–retest reliability than cortical surface area or volume^44,45^. These characteristics may contribute to the lower replicability observed for cortical thickness in the present study.

The cross-omics analyses further showed that replicability differed markedly across molecular layers. In matched participants, proteomic associations had higher KDRD estimates than pQTL associations across all 45 evaluated traits, whereas the relative replicability of proteomic and metabolomic associations varied across traits, with metabolomics showing higher estimates for a larger proportion of traits. These findings indicate that association replicability is not uniform across different types of molecular data. The observed cross-omics differences may arise from both biological and statistical sources. Proteins and metabolites are more proximal to phenotypic states than genetic variation and may therefore capture biological processes more directly related to the traits^4,41,46^. In addition, the metabolomic analyses involved substantially fewer tested features than the proteomic analyses (249 metabolites versus 2,920 proteins), resulting in a less stringent multiple-testing threshold. Differences in the number of tested features and the corresponding significance thresholds may therefore also contribute to the observed variations in replicability across molecular layers.

Data quality was also closely related to proteomic discovery replicability. Higher protein missingness, a greater proportion of samples below the limit of detection, a higher proportion of low-abundance proteins and higher protein-level assay variability (CoV) were consistently associated with lower KDRD. These characteristics can reduce the precision and stability of measured protein levels and consequently affect the association statistics on which significant discoveries are based^4,38^. The observed relationships therefore emphasize that replicability in large-scale proteomic association studies depends not only on sample size but also on the quality of the underlying molecular measurements. These findings are also important for the interpretation of KDRD assessments. KDRD assesses expected replicability under comparable experimental and analytical conditions, whereas our results indicate that proteomic data-quality characteristics can substantially influence replicability. If a future proteomic panel has poorer data quality than the dataset used for KDRD assessment, the KDRD value may overestimate the replicability that would actually be observed.

KDRD can also be used to estimate replicability at target sample sizes larger than that of the observed dataset. This capability is relevant to large-scale studies in which molecular profiling is being extended to larger cohorts. For example, proteomic profiling in the UK Biobank is being extended from approximately 54,000 participants to the full cohort of approximately 500,000 participants^47^. Our sample-size evaluation showed that KDRD was projected to increase with sample size, although the magnitude of improvement varied across traits. At a target sample size of approximately 200,000, all 45 evaluated traits were projected to achieve at least moderate replicability, whereas a sample size of 500,000 was projected to achieve high replicability across all traits. Notably, the planned full-cohort sample size of the UK Biobank reaches the level at which high replicability was estimated for all evaluated traits. These estimates assume that the underlying association effects and experimental and analytical conditions remain comparable as the sample size increases.

Several limitations are important when interpreting KDRD. First, KDRD estimates replicability for a hypothetical independent study conducted under comparable experimental and analytical conditions. Differences in population characteristics, phenotype definitions, molecular measurements, data processing or statistical procedures between studies can introduce additional heterogeneity that is not represented by the KDRD assessment^28,30,31^. KDRD therefore provides an assessment of replicability conditional on comparable study conditions and should not be regarded as a substitute for external validation when the aim is to evaluate generalizability across heterogeneous populations or study settings. Second, the sample-size adjustment assumes that the estimated proportion of null associations remains unchanged across different target sample sizes. However, the estimated value of this proportion may change with sample size, which may affect the accuracy of KDRD-based sample-size evaluation. Estimates for target sample sizes far beyond that of the observed dataset should therefore be considered exploratory rather than definitive. In addition, when the reference dataset used for KDRD assessment has a relatively small sample size, some model parameters may be estimated less accurately, including the proportion of null associations. This uncertainty may further affect the results of KDRD assessments.

In summary, KDRD provides a framework for estimating the replicability of statistically significant discoveries based on a single set of association statistics. Across forty-five brain-related traits, KDRD showed strong agreement with DSC values observed between paired random subsamples, enabling replicability assessment without splitting the original sample. Its applications further revealed substantial variations in proteomic discovery replicability across traits and molecular layers, identified associations between proteomic data quality and replicability, and enabled evaluation of replicability at target sample sizes beyond those currently available. Together, these findings demonstrate that KDRD provides a practical and flexible framework for quantitative replicability assessment in large-scale association studies. It is particularly useful for studies based on a single available dataset. KDRD-based sample-size evaluation can further estimate the sample size required to achieve desired replicability, including sample sizes beyond those currently available.

## Data availability

The UK Biobank data used in this study included proteomic, genomic, metabolomic, brain imaging, and phenotypic data. These data were accessed under UK Biobank application number 19542 and are subject to controlled access under UK Biobank governance and data-protection requirements; therefore, they cannot be redistributed by the authors. Access should be requested directly from UK Biobank through its Access Management System by eligible researchers, subject to UK Biobank approval and its data-use agreement. Access enquiries can be directed to the UK Biobank Access Management Team (https://www.ukbiobank.ac.uk/). Under the standard UK Biobank access procedure, submitted applications are escalated for review within 10 working days, with feedback provided thereafter. The pQTLs used for the multi-omics comparison were obtained from the pQTL results reported in the supplementary materials of a previously published study^38,40^. Additional non-controlled data generated in this study are available from the corresponding author (i.e. Jianfeng Feng) upon reasonable request.

## Code availability

The code used in this study is available from the corresponding author, Jianfeng Feng, upon reasonable request. Analyses were performed using R version 4.3.0.

## Acknowledgements

We thank Dr. Xiuqin Liu from the University of Science and Technology Beijing for her helpful discussions and valuable suggestions.

## Author contributions

Conception: Zeyu Jiao, Yinglei Lai, Yixin Zhang and Jianfeng Feng. Design: Zeyu Jiao, Yixin Zhang, Yinglei Lai and Jianfeng Feng. Data acquisition, processing and analysis: Zeyu Jiao, Yixin Zhang, Yinglei Lai, Yawen Yang, Jujiao Kang, Wei Zhao and Jianfeng Feng. Manuscript writing and revising: Zeyu Jiao, Yixin Zhang, Yinglei Lai, Jujiao Kang, Liang Ma and Jianfeng Feng.

## Extended Data

**Extended Data Figure 1.**
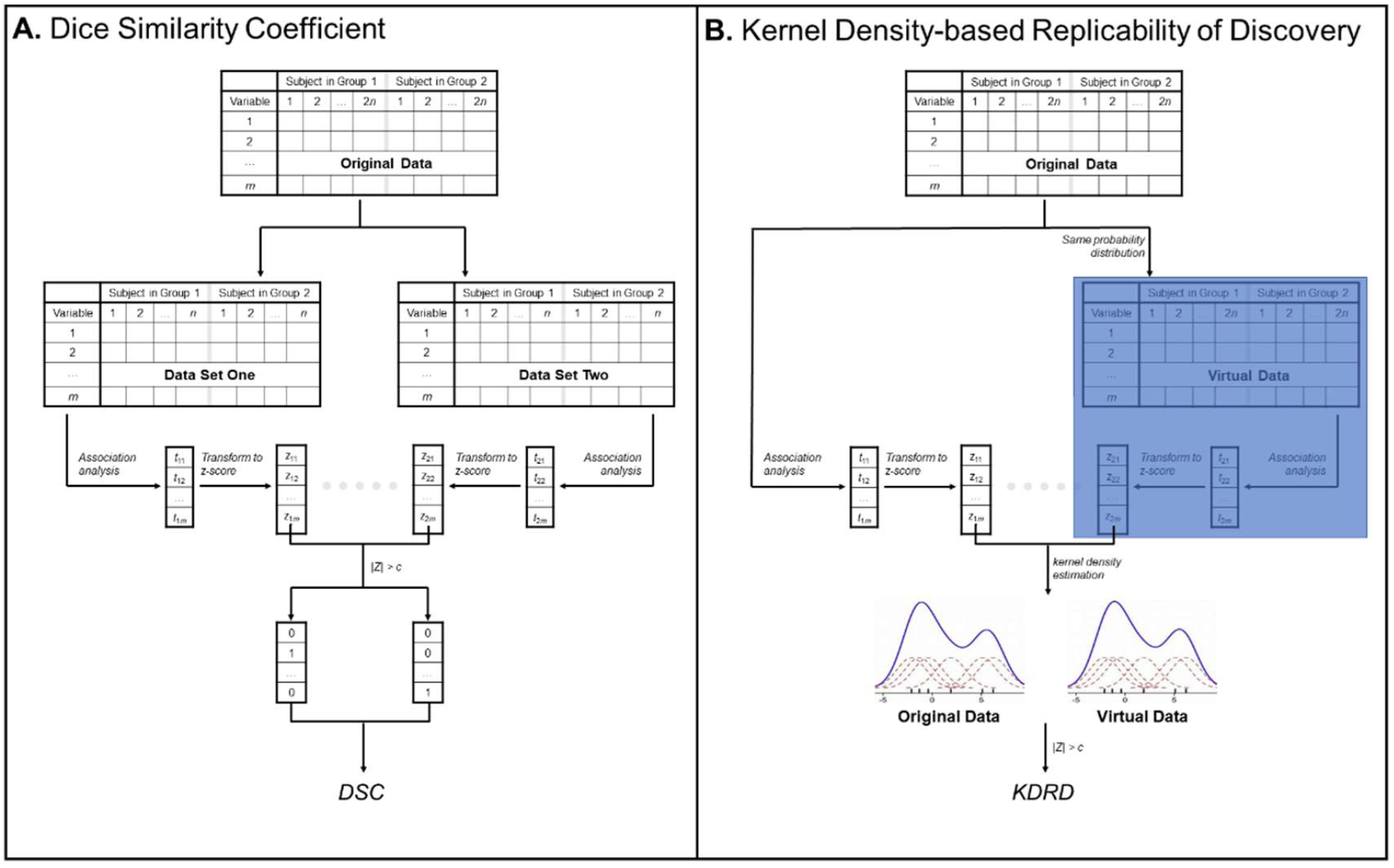
Overview of the analytical steps to estimate replicability in this study. **A**. the analytical steps for *DSC*. **B**. the analytical steps for *KDRD*.

## References

1. National Academies of Sciences, Engineering, and Medicine. Reproducibility and Replicability in Science (National Academies Press, 2019).

2. Goodman, S. N., Fanelli, D. & Ioannidis, J. P. A. What does research reproducibility mean? Sci. Transl. Med. 8, 341ps12 (2016).

3. Ioannidis, J. P. A., Thomas, G. & Daly, M. J. Validating, augmenting and refining genome-wide association signals. Nat. Rev. Genet. 10, 318–329 (2009).

4. Eldjarn, G. H. et al. Large-scale plasma proteomics comparisons through genetics and disease associations. Nature 622, 348–358 (2023).

5. Draisma, H. H. M. et al. Genome-wide association study identifies novel genetic variants contributing to variation in blood metabolite levels. Nat. Commun. 6, 7208 (2015).

6. Marek, S. et al. Reproducible brain-wide association studies require thousands of individuals. Nature 603, 654–660 (2022).

7. Storey, J. D. & Tibshirani, R. Statistical significance for genomewide studies. Proc. Natl Acad. Sci. USA 100, 9440–9445 (2003).

8. Efron, B. Correlated z-values and the accuracy of large-scale statistical estimates. J. Am. Stat. Assoc. 105, 1042–1055 (2010).

9. Heller, R., Bogomolov, M. & Benjamini, Y. Deciding whether follow-up studies have replicated findings in a preliminary large-scale omics study. Proc. Natl Acad. Sci. USA 111, 16262–16267 (2014).

10. McGuire, D. et al. Model-based assessment of replicability for genome-wide association meta-analysis. Nat. Commun. 12, 1964 (2021).

11. Jiao, Z. et al. A model-based approach to assess reproducibility for large-scale high-throughput MRI-based studies. NeuroImage 255, 119166 (2022).

12. Li, Q., Brown, J. B., Huang, H. & Bickel, P. J. Measuring reproducibility of high-throughput experiments. Ann. Appl. Stat. 5, 1752–1779 (2011).

13. Zhao, Y., Sampson, M. G. & Wen, X. Quantify and control reproducibility in high-throughput experiments. Nat. Methods 17, 1207–1213 (2020).

14. Maitra, R. A re-defined and generalized percent-overlap-of-activation measure for studies of fMRI reproducibility and its use in identifying outlier activation maps. NeuroImage 50, 124–135 (2010).

15. Wang, J., Liang, H., Zhang, Q. & Ma, S. Replicability in cancer omics data analysis: measures and empirical explorations. Brief. Bioinform. 23, bbac304 (2022).

16. Turner, B. O., Paul, E. J., Miller, M. B. & Barbey, A. K. Small sample sizes reduce the replicability of task-based fMRI studies. Commun. Biol. 1, 62 (2018).

17. Kang, G. et al. Evaluation of a two-step iterative resampling procedure for internal validation of genome-wide association studies. J. Hum. Genet. 60, 729–738 (2015).

18. Sabourin, J. A. et al. ComPaSS-GWAS: A method to reduce type I error in genome-wide association studies when replication data are not available. Genet. Epidemiol. 43, 102–111 (2019).

19. Ioannidis, J. P. A. Why most discovered true associations are inflated. Epidemiology 19, 640–648 (2008).

20. Button, K. S. et al. Power failure: why small sample size undermines the reliability of neuroscience. Nat. Rev. Neurosci. 14, 365–376 (2013).

21. Palmer, C. & Pe’er, I. Statistical correction of the Winner’s Curse explains replication variability in quantitative trait genome-wide association studies. PLoS Genet. 13, e1006916 (2017).

22. Greenbaum, D., Sboner, A., Mu, X. J. & Gerstein, M. Genomics and privacy: implications of the new reality of closed data for the field. PLoS Comput. Biol. 7, e1002278 (2011).

23. Byrd, J. B. et al. Responsible, practical genomic data sharing that accelerates research. Nat. Rev. Genet. 21, 615–629 (2020).

24. Kaye, J. The tension between data sharing and the protection of privacy in genomics research. Annu. Rev. Genomics Hum. Genet. 13, 415–431 (2012).

25. Bonomi, L., Huang, Y. & Ohno-Machado, L. Privacy challenges and research opportunities for genomic data sharing. Nat. Genet. 52, 646–654 (2020).

26. Kässens, J. C., Wienbrandt, L. & Ellinghaus, D. BIGwas: single-command quality control and association testing for multi-cohort and biobank-scale GWAS/PheWAS data. GigaScience 10, giab047 (2021).

27. Schönherr, S. et al. Performing highly parallelized and reproducible GWAS analysis on biobank-scale data. NAR Genom. Bioinform. 6, qae015 (2024).

28. Roberts, G. H. L. et al. Expanded COVID-19 phenotype definitions reveal distinct patterns of genetic association and protective effects. Nat. Genet. 54, 374–381 (2022).

29. Dahl, A. et al. Phenotype integration improves power and preserves specificity in biobank-based genetic studies of major depressive disorder. Nat. Genet. 55, 2082–2093 (2023).

30. Botvinik-Nezer, R. et al. Variability in the analysis of a single neuroimaging dataset by many teams. Nature 582, 84–88 (2020).

31. Gatidis, S. et al. Better together: data harmonization and cross-study analysis of abdominal MRI data from UK Biobank and the German National Cohort. Invest. Radiol. 58, 346–354 (2023).

32. Bastarache, L. et al. The phenotype-genotype reference map: Improving biobank data science through replication. Am. J. Hum. Genet. 110, 1522–1533 (2023).

33. Wasfi, R. et al. Recruiting participants for population health intervention research: effectiveness and costs of recruitment methods for a cohort study. J. Med. Internet Res. 23, e21142 (2021).

34. Milbourn, H. et al. Effectiveness and costs of participant recruitment strategies to a web-based population cohort: observational study. J. Med. Internet Res. 27, e75116 (2025).

35. Liu, S. et al. Replicable brain–phenotype associations require large-scale neuroimaging data. Nat. Hum. Behav. 7, 1344–1356 (2023).

36. Kang, K. et al. Study design features increase replicability in brain-wide association studies. Nature 636, 719–727 (2024).

37. Sudlow, C. et al. UK Biobank: An open access resource for identifying the causes of a wide range of complex diseases of middle and old age. PLoS Med. 12, e1001779 (2015).

38. Sun, B. B. et al. Plasma proteomic associations with genetics and health in the UK Biobank. Nature 622, 329–338 (2023).

39. Bycroft, C. et al. The UK Biobank resource with deep phenotyping and genomic data. Nature 562, 203–209 (2018).

40. Jiao, Z. et al. Merits and challenges of plasma proteomics on association replicability. bioRxiv 2025.09.01.673490 (2025).

41. Julkunen, H. et al. Atlas of plasma NMR biomarkers for health and disease in 118,461 individuals from the UK Biobank. Nat. Commun. 14, 604 (2023).

42. Alfaro-Almagro, F. et al. Image processing and quality control for the first 10,000 brain imaging datasets from UK Biobank. NeuroImage 166, 400–424 (2018).

43. Davies, G. et al. Study of 300,486 individuals identifies 148 independent genetic loci influencing general cognitive function. Nat. Commun. 9, 2098 (2018).

44. Grasby, K. L. et al. The genetic architecture of the human cerebral cortex. Science 367, eaay6690 (2020).

45. Hedges, E. P. et al. Reliability of structural MRI measurements: The effects of scan session, head tilt, inter-scan interval, acquisition sequence, FreeSurfer version and processing stream. NeuroImage 246, 118751 (2022).

46. IGVF Consortium. Deciphering the impact of genomic variation on function.Nature 633, 47–57 (2024).

47. Largest proteome study enlists 14 biopharmas. Nat. Biotechnol. 43, 153 (2025).

